# Suppression of glycolysis decreases sugar-induced cell death in Saccharomyces cerevisiae

**DOI:** 10.1101/2024.08.17.608408

**Authors:** Airat Ya. Valiakhmetov

**Affiliations:** Laboratory of regulation of biochemical processes, Skryabin Institute of Biochemistry and Physiology of Microorganisms, Federal Research Center “Pushchino Scientific Center for Biological Research of the Russian Academy of Sciences”, Pushchino 142290, Russia

**Keywords:** yeast, trehalose, sugar-induced cell death, glycolysis, necrosis, pentose-phosphate pathway

## Abstract

Although 30 years have passed since the description of sugar-induced cell death (SICD), the specific molecular mechanism that triggers this process remains unclear. This paper attempts to shed light on the relationship between SICD and glucose catabolism. Deletion of the TPS1 gene resulted in a 44% suppression of SICD and a 75% reduction in the number of cells with excess ROS. The suppression was comparable to the suppression of SICD (38%) and ROS (71%) with deletion of the HXK2 gene. Since HXK2 is the first enzyme in the glycolytic pathway, the effect of two other key glycolytic enzymes on SICD was tested. Deletion of the TDH3 gene (glyceraldehyde-3-phosphate dehydrogenase) resulted in a 39% suppression of SICD and ROS by 48%. Inhibition of Tdh3p with 1 mM iodoacetamide also suppressed SICD by 67% and ROS by 58%. Deletion of the PFK1 (phosphofructokinase 1) gene resulted in a complete block of SICD (97%), but unexpectedly resulted in a significant increase in the number of cells with excess ROS. All strains tested (except ΔPFK1) showed increased glucose consumption, suggesting that redistribution of glucose fluxes between glycolysis and the pentose phosphate shunt is a key regulator of SICD development.

## Introduction

In 1991, an interesting observation was described: the death of yeast cells during incubation with glucose alone in the absence of other nutrients (Granot and Snyder 1991). This phenomenon is called SICD (**s**ugar-**i**nduced **c**ell **d**eath) (Granot and Dai 1997). SICD in yeast in the stationary phase has of apoptotic nature and is caused by ROS accumulation (Granot et al. 2003). Phosphorylation of glucose or fructose was a prerequisite for SICD (Granot and Snyder 1993; Granot and Dai 1997). SICD did not depend on the glucose high-affinity transport system SNF3 and was caused by a wide range of sugars (Granot and Snyder 1993). It’s worth noting that SICD has been observed in the bottom-fermenting yeast *Saccharomyces pastorianus* under strictly anaerobic conditions as well (Yoshimoto et al. 2009). In attempts to elucidate the specific mechanism underlying SICD, attention was paid to the Crabtree effect (Lee et al. 2011; de Alteriis et al. 2018). The Crabtree effect is hypothesized to involve competition between the mitochondrial respiratory chain and glycolytic enzymes for ADP and inorganic phosphate. In the authors’ opinion, the inhibition of SICD in stationary yeast cultures by exogenous phosphate and succinate confirms this hypothesis. It is important to note here that all of these studies were carried out with yeast in the stationary growth phase. Later it was shown that incubation of exponentially growing *S. cerevisiae* yeast exposed to glucose in the absence of other necessary nutrients for growth also led to SICD, but it displayed characteristics of primary necrosis. In contrast to stationary cells, SICD in exponentially growing yeast occurred much more rapidly (minutes compared to days in stationary-phase cells) as a result of ruptures of nuclear, mitochondrial, and plasma membranes and affected only cells in the S phase of the cell cycle (Valiakhmetov et al. 2019). A remarkable characteristic of SICD observed in the mid-log-phase culture was its responsiveness to extracellular pH and membrane potential. Incubation of yeast with glucose at pH 7.0 (dissipation ΔpH) or in the presence of 150 mM KCl (dissipation ΔΨ) completely prevented SICD (Bidiuk et al. 2021). These facts raise doubts about the idea that primary necrosis occurs only due to extreme external physical or chemical impact on the cell. SICD occurs when the cell is not exposed to any extreme external impacts. Instead, an imbalance in the availability of all nutrients required for growth resulted in SICD through primary necrosis. Notably, the mitochondrial respiratory chain is not involved in the development of SICD in exponentially growing yeast. This was demonstrated in both the case of petite mutants obtained by treating yeast cells with ethidium bromide and in the ΔAFO1 strain, which lacks a large mitochondrial ribosomal subunit (Bidiuk et al. 2021).

Trehalose is one of two storage carbohydrates in yeast (François and Parrou 2001). The biosynthesis of trehalose and its relationship with cellular signaling pathways has been the subject of numerous studies (Gancedo and Flores 2004; Paul et al. 2008). Of particular interest is the proposed link between glucose entry into glycolysis and the activity of a trehalose-6-phosphate synthetase (EC 2.4.1.15 Tps1p) (Neves et al. 1995; Thevelein and Hohmann 1995; Hohmann et al. 1996; van Heerden et al. 2014; Van Leemputte et al. 2020).

Additionally, TPS1 deletion resulted in multiple pleiotropic effects (Pereira et al. 2001). The most striking of these is the inability of the ΔTPS1 strain to utilize glucose for growth (Hottiger et al. 1987; Hohmann et al. 1993; Hohmann et al. 1994). Deletion of the TPS1 gene results in altered thermosensitivity and protein folding defects (Singer and Lindquist 1998b; Singer and Lindquist 1998a), confers desiccation tolerance (Tapia et al. 2015), and reduces sporulation efficiency (De Silva-Udawatta and Cannon 2001; Liu et al. 2020).

Due to such a wide range of effects of trehalose and Tps1p on the metabolism of yeast cells, the task arose to test the effect of Tps1p on SICD.

## 2. Materials and methods

### 2.1 Culture growth

Strains ΔTPS1, ΔTPS2, ΔHXK2, ΔTDH3, ΔPFK1 and their parental strain BY4742 were obtained from the Euroscarf collection. The culture grew on the standard YPD medium (Applichem, Darmstadt, Germany) for 15– 17 h (mid-exponential phase) and was twice washed with distilled water. Yeast cells were pelleted and suspended (1 g/10 mL w/w, 1.45 × 10^8^ cells/mL) in MilliQ water.

### 2.2 1,2,3-Dihydrorhodamine (DHR) Staining

To estimate the number of cells with elevated intracellular ROS levels, we used DHR (Sigma-Aldrich, St. Louis, MO, USA). 5 µL of 0.5 mg/mL DHR in DMSO was added to 250 µL cell suspension in water. The samples were incubated in a ThermoMixer (Eppendorf, Hamburg, Germany) for 1 h at 30°. The cells were pelleted at 13,000× g for 1 min, and the supernatant was discarded. The cells were washed once with 0.5 mL of water, and the cell pellet was suspended in 250 µL water. Another 250 µL of cell suspension in water without the addition of DHR was subjected to all the above steps to obtain cells for propidium iodide (PI) staining (see below).

### 2.3 SICD Assay and Flow Cytometry

SICD assay for DHR-stained and unstained cells was done in a parallel series of tubes. 250 µL of cells obtained as described above was distributed in 50 µL into four tubes. This was the standard test kit for the SICD. Nothing was added to the first tube—this was the control. The second tube served as the main sample for checking the SICD. In the third tube, water was replaced with buffer (50 mM HEPES, pH 7.0) to test the effects of neutral pH on the SICD. 2.5 µL of 3 M KCl was added to the 4th tube to check the value of the membrane potential on SICD. Then, to induce SICD, 2.5 µL of 2 M glucose solution was added to all tubes (final [C] = 100 mM), except for the 1st one. The samples were incubated in a ThermoMixer (Eppendorf, Hamburg, Germany) at 30 °C for 1 h. To proceed to flow cytometry to the samples with DHR-stained cells 1 mL of water was added. PI staining was used to determine the percentage of dead cells. To the samples with not DHR-stained cells, 150 µL of 4 µg/mL PI (Sigma-Aldrich, St. Louis, MO, USA) was added, and after brief (1–2 min) incubation at RT, 0.8 mL of water was added. All samples were kept on ice. A total of 100,000 cells were counted at each experimental point using a NovoCyte Flow cytometer (Agilent, Santa Clara, CA, USA). DHR-stained cells were detected using 488 nm for excitation and 530/30 nm for emission. PI-stained cells were detected using 488 nm for excitation and 572/28 nm for emission. All assays were repeated three times, and the mean results are presented.

### 2.4 Iodoacetamide (IAD) assay

Cells, prepared as in *2.2* were distributed in 50 μL into 7 tubes. 1^st^ tube – control contained no any additions. The 2^nd^ tube served as the main sample for checking the SICD. One μL of the stock solutions of IAD (12.5, 25, 37.5, 50, and 62.5 mM in water) was added to tubes 3-7, giving final concentrations of IAD 0.25, 0.5, 0.75, 1, and 1.25 mM respectively. After 20 min incubation at 30°C, to induce SICD 2.5 µL of 2 M glucose solution was added to all tubes (final [C] = 100 mM), except for the 1^st^ one. The samples were incubated at 30 °C for 1 h. Then samples proceed to flow cytometry as in *2.3*.

### 2.5 Glucose uptake assay and pH measurement

15 ml of cell suspension prepared as in section *2.1* were placed in a thermostated cell (30°C) with a magnetic stirrer. After temperature stabilization 0.75 ml of 2 M glucose was added. 100 μL of the cell suspension was immediately taken and spun down at 18 000 x g (starting point). Super was transferred into a clean tube. Other samples were taken every 10 minutes the same way and kept on ice. To assess the glucose concentration, a colorimetric method based on GOD-PAP was used using the “Glucose DDC” kit (Diakon-DC, Pushchino, Russia) according to the instructions. To measure pH, a pH electrode and temperature sensor connected to a pH meter were immersed in the same 15 ml suspension in which the glucose concentration was measured.

### 2.6 Statistical analysis

All measurements were carried out at least 3 times. Data are presented as mean ± SD. One-way ANOVA statistics were calculated using the Prism 9 software (GraphPad, Boston, MA, USA).

## Results and discussion

Since the enzymes involved in trehalose biosynthesis show pleiotropic effects, we tested the effect of deletion of the TPS1 and TPS2 genes on SICD and ROS production when *S. cerevisiae* cells were incubated with 100 mM glucose alone in the absence of other nutrients. There is an important note to make here. All studies related to the study of the yeast TPS1/TPS2 complex note that these deletion strains do not grow on fermentable sugars, in particular glucose. However, several studies have been published in which the growth of ΔTPS1 cells on glucose was observed and this growth was stimulated by a peptone-rich medium (Walther et al. 2013; van Heerden et al. 2014; Gibney et al. 2020; Chen et al. 2022). At this point, it has been proposed that the genetic background of the ΔTPS1 strains plays a key role in explaining the observed contradiction (Neves et al. 1995; Walther et al. 2013; Chen et al. 2022). In this study, ΔTPS1 is derived from the parental strain BY4742. After plating from frozen stock onto a YPD plate, the cells begin to grow with a delay of 36-48 hours and then continue to grow as the parent strain. A similar effect was previously observed on the ΔTPS1 strain derived from the haploid strain *S. cerevisiae* CEN.PK133-5D ura3-52 (Walther et al. 2013). After the subculture of our cells (BY4742:ΔTPS1) from solid to liquid YPD, their growth rate does not differ from the growth rate of the parental strain BY4742.

As shown in Fig. 1, deletion of the TPS2 gene did not lead to any significant change in SICD and ROS levels. This can be explained as follows. Trehalose-6-phosphate phosphatase (EC 3.1.3.12 Tps2p) catalyzes the conversion of trehalose-6-phosphate (T-6-P) to trehalose. However, T-6-R begins to accumulate in the cell when available glucose in the growth medium is depleted or stress-forming conditions appear (Lillie and Pringle 1980; Hottiger et al. 1987; Parrou et al. 1997). Under our conditions, the cell culture is in the middle of the log phase of growth and optimal conditions for growth, and therefore there should not be any significant reserves of T-6-P for Tps2p activity. At the same time, in the ΔTPS1 strain, SICD was suppressed by 44%, and the number of cells with excess ROS was reduced by 75%. Since glucose phosphorylation is a prerequisite for SICD (Granot and Snyder 1993; Granot and Dai 1997), it is logical to hypothesize a possible link between SICD, glycolysis, and Tps1p activity. The current main hypothesis linking trehalose biosynthesis with glycolysis suggests that T-6-P (or Tps1p itself) inhibits hexokinase 2 (EC 2.7.1.1 Hxk2p) and thereby blocks the entry of glucose into glycolysis by redirecting the flow of glucose to the trehalose biosynthesis pathway (Thevelein and Hohmann 1995; Hohmann et al. 1996; Bonini et al. 2003). Therefore, it would be logical to accept the hypothesis of Hxk2p inhibition by T-6-P or Tps1p to explain our data. However, in 2023, the proteome epistatic database (https://y5k.bio.ed.ac.uk) became available (Messner et al. 2023) which revealed a very interesting detail. In the ΔTPS1 strain, the expression of Hxk2p is almost 5-fold reduced (Supplemental Fig. 1). This should lead to a significant reduction in the amount of phosphorylated glucose and, consequently, to the suppression of SICD. To test this hypothesis, the effect of HXK2 gene deletion on SICD and ROS was examined. As shown in Fig. 1, in the ΔHXK2 strain, the levels of SICD and ROS decreased to the same extent as in the ΔTPS1 strain (SICD by 38%, the number of cells with excess ROS by 71%). Based on these data, it can be assumed that the suppression of SICD and ROS generation in the ΔTPS1 strain is the result of a drop in the amount of phosphorylated glucose as a result of significantly reduced expression of Hxk2p. This fact prompted us to test the effect on SICD and ROS generation of two more key glycolytic enzymes—6-phosphofructokinase (EC 2.7.1.11 Pfk1p), encoded by the PFK1 gene, and glyceraldehyde-3-phosphate dehydrogenase (EC 1.2.1.12 Tdh3p), encoded by the TDH3 gene. Deletion of TDH3 resulted in a 40% suppression of SICD and a 48% reduction in the number of cells with excess ROS (Fig. 1). Inhibition of Tdh3p by iodoacetamide also resulted in suppression of SICD and ROS (Fig. 2).

**Fig. 1.**
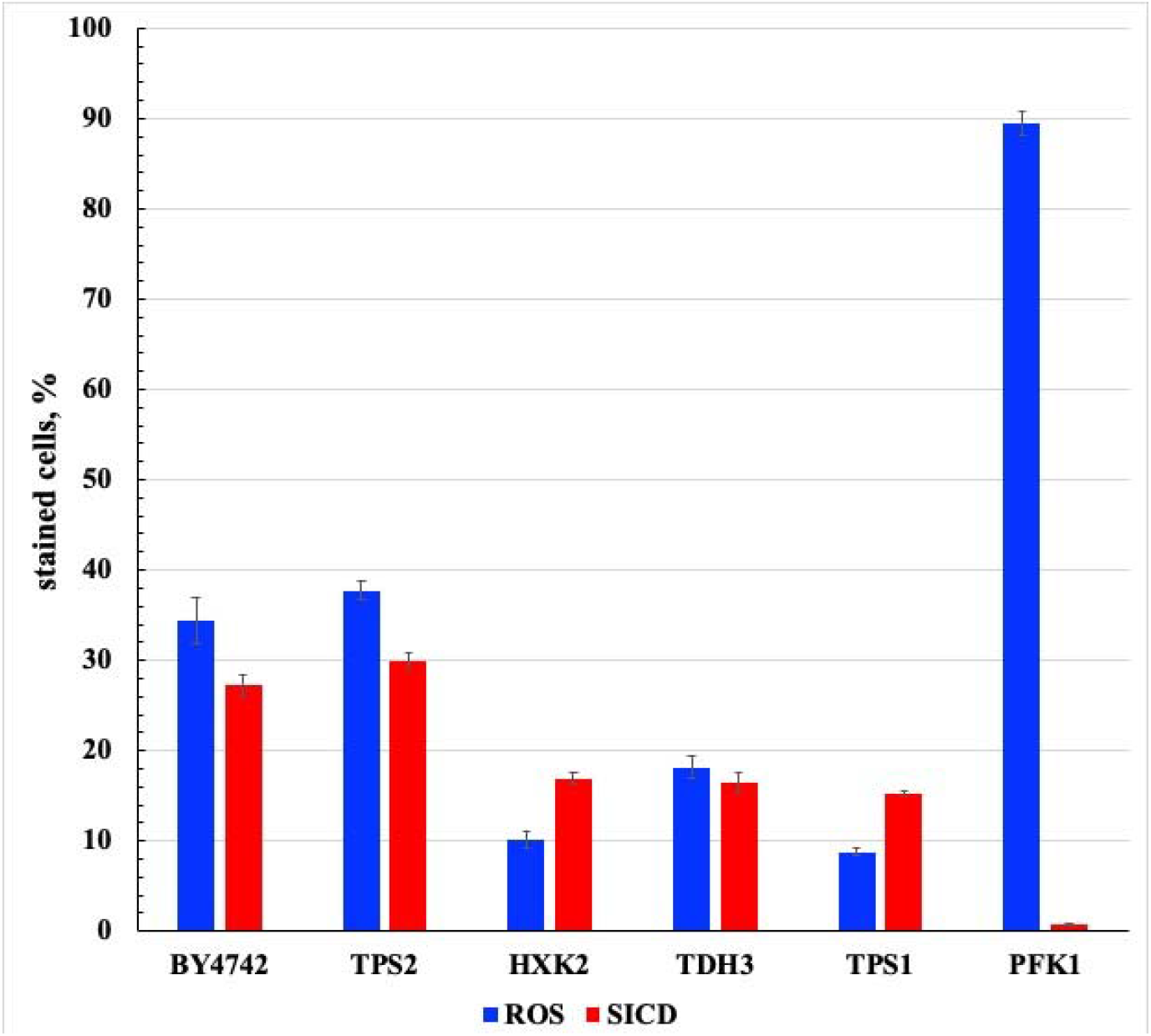
SICD and ROS generation in BY4742 and in deletion strains. The data represents the percentage of stained cells out of the total number of cells counted (100,000 cells). Cells were incubated in MilliQ water with 100 mM glucose at 30°C for 1 hour. ROS were detected by DHR fluorescence. PI fluorescence was used to detect dead cells (SICD). Individual strains were compared to parental strain BY4742 using a one-way ANOVA test. P-value for all strains (except ΔTPS2) was < 0.0001. P-value for BY4742 vs. ΔTPS2 was >0.1 for SICD and ROS. Data are presented as mean±SD from 3 independent experiments.

**Fig. 2.**
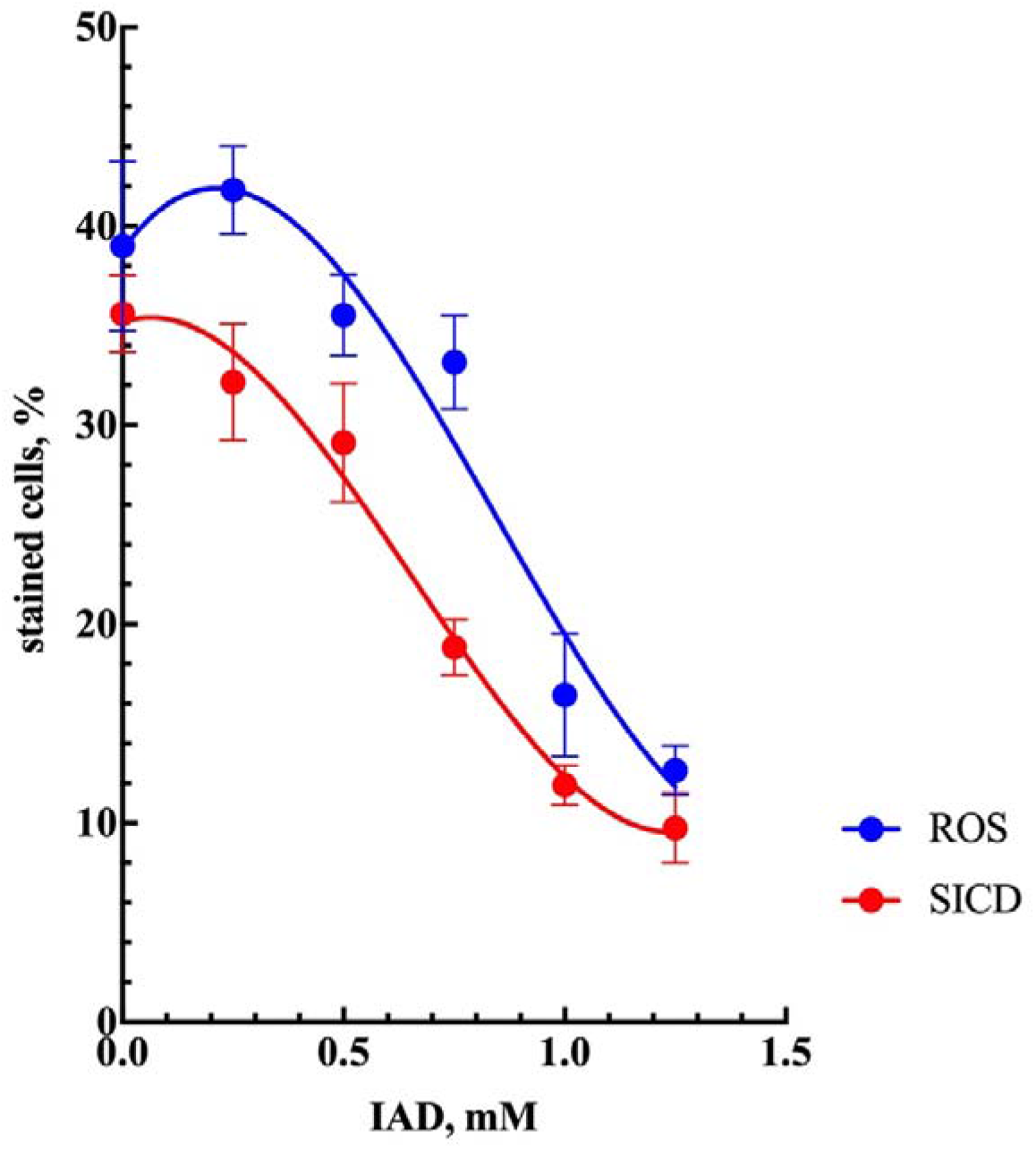
Effect of Tdh3p (glyceraldehyde-3-phosphate dehydrogenase) inhibitor IAD on SICD and ROS generation in strain BY4742. Data represent the percentage of stained cells out of the total number of cells counted (100,000 cells). Cells were incubated with the indicated concentrations of IAD for 20 min. After this, glucose was added at a final concentration of 100 mM and incubated for 1 hour at 30°C. ROS were detected by DHR fluorescence. PI fluorescence was used for dead cell detection (SICD). Data are presented as mean±SD from 3 independent experiments.

The putative mechanism of the effect of Tdh3p on SICD will be discussed below. Deletion of the PFK1 gene resulted in complete suppression of SICD and, unexpectedly, a very significant increase in the number of cells with ROS (Fig. 1). It is important to note that the fluorescent probe for ROS, 1,2,3-DHR, can be oxidized to rhodamine by a wide range of ROS (Kalyanaraman et al. 2012). Along with H_2_O_2_, which also serves as part of the cellular signaling system (D’Autréaux and Toledano 2007), other spices of ROS (superoxide, singlet oxygen, and the hydroxyl radical) can be produced in the cell (Murphy et al. 2011). As has been shown previously, SICD occurs due to rapid disruption of the integrity of plasma, mitochondrial, and vacuolar membranes (Valiakhmetov et al. 2019). Such a destructive effect can most likely only be exerted by the short-lived hydroxyl radical. Therefore, we believe that the increase in the number of cells with ROS observed in the ΔPFK1 strain reflects the accumulation of relatively harmless H_2_O_2_, which does not lead to cell death. Pfk1p, along with pyruvate kinase (EC 2.7.1.40), is an enzyme that works in glycolysis only in the forward direction and its deletion leads to a block in glycolysis (Lobo and Maitra 1983). Under our conditions, this is confirmed by the virtual absence of acidification of the medium during incubation of ΔPFK1 with 100 mM glucose and extremely low glucose consumption compared to the parental strain BY4742 (Fig. 3).

**Fig. 3.**
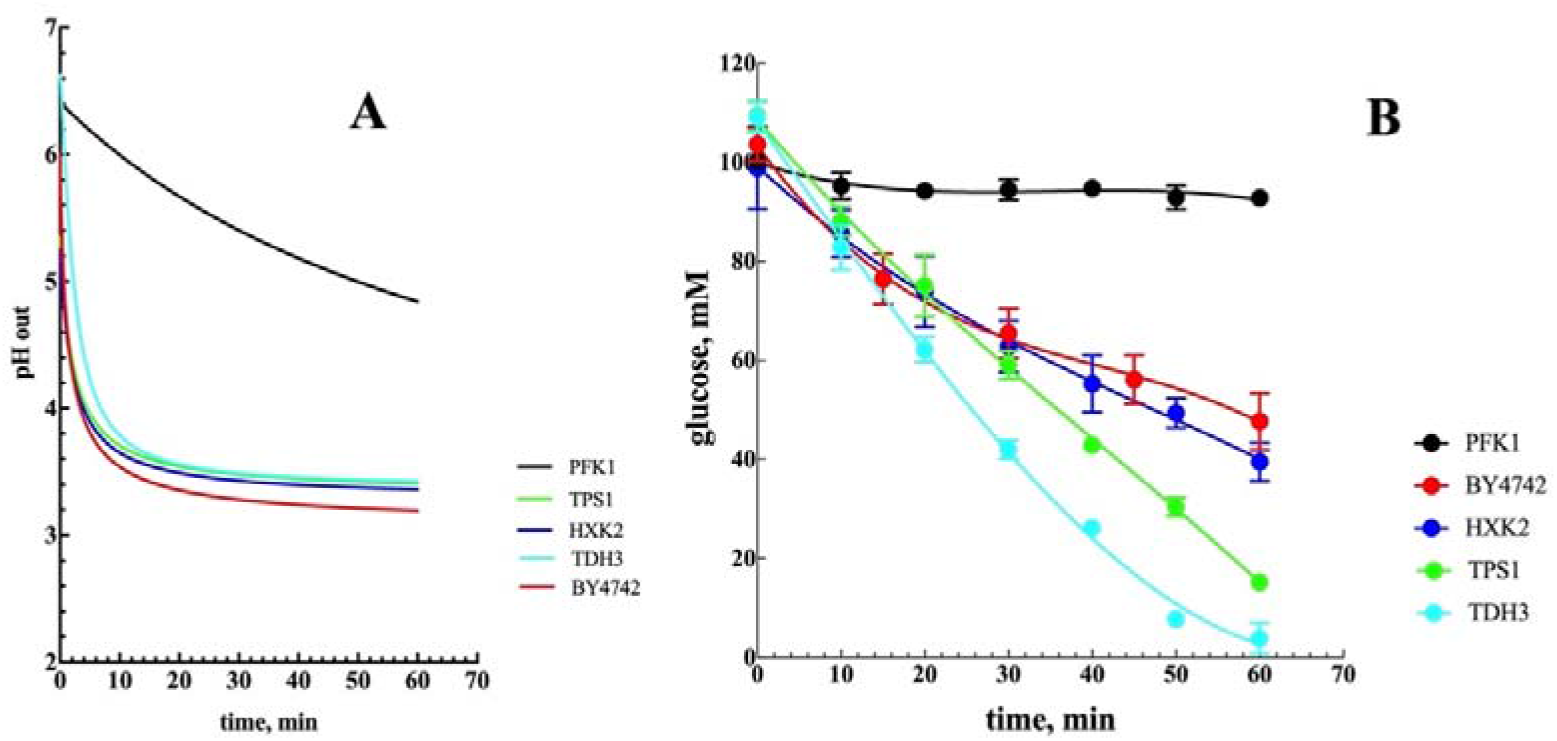
**A** - Dynamics of pH of the medium during incubation of BY4742 cells and deletion strains with 100 mM glucose at 30°C. **B** – Changes in the concentration of glucose in the medium when cells are incubated with 100 mM glucose at 30°C. The data presented in panels A and B were obtained simultaneously from the same experiment. In panel **B** data are presented as mean±SD from 3 independent experiments.

The suppression of SICD and ROS in all tested strains suggests that the full functioning of glycolysis is the main condition for the development of SICD. Partial (ΔTPS1, ΔHXK2, ΔTDH3, iodoacetamide) or complete (ΔPFK1) suppression of glycolysis resulted in suppression of SICD. However, the question arises - what is the fate of glucose, the consumption of which in all strains (except ΔPFK1) exceeds the level of consumption in the parent strain BY4742 (Fig. 3B). There are two main pathways for glucose catabolism in yeast - glycolysis and the pentose phosphate shunt (PPS). A special feature of PPS is the supply of cells with NADPH, which is used, in particular, to restore oxidized glutathione, and the production of ribulose-5-phosphate, which is then involved in the synthesis of RNA and DNA. In exponentially growing yeast, SICD is observed in cells in the S-phase of the cell cycle, when the functioning of the PPS is most required by the cell (Valiakhmetov et al. 2019). We hypothesize the following chain of events leading to the development of SICD. When cells are incubated with glucose in the absence of other nutrients, glucose does not enter the PPS in cells in the S-phase for reasons that are still unclear. This leads to a deficiency of NADPH and therefore an inability to reduce oxidized glutathione. As a result, there is an excessive accumulation of hydroxyl radicals, which in turn leads to the oxidation of membrane lipids with their subsequent rupture. However, it appears that it is possible to partially restore PPS function by suppressing glycolysis. In ΔTPS1 and ΔHXK2 strains, glucose can be phosphorylated by hexokinase 1 (EC 2.7.1.1 Hxk1p) and redirected to PPS. This hypothesis is supported by the fact that TPS1 is involved in the regulation of PPS in the pathogenic filamentous fungus *Magnaporthe grisea* (Wilson et al. 2007). Excess glyceraldehyde-3-phosphate (in the ΔTDH3 strain and when Tdh3p is inhibited by iodoacetamide) can also enter the PPS, leading to NADPH production and partial reduction of oxidized glutathione followed by hydroxyl radical detoxification (SICD inhibition).

The observed acidification of the medium during incubation of all (except ΔPFK1) strains with glucose remains unclear (Fig. 3A). Obviously, this is not a consequence of the release of protons during the hydrolysis of ATP of glycolytic origin by H^+^-ATPase of the plasma membrane because glycolysis (at least at its payoff stage) is suppressed. Most likely, this may be a consequence of the hydrolysis of ATP produced during the functioning of the respiratory chain in mitochondria (all our experiments were carried out under aerobic conditions), and/or a consequence of the release of octanoic acid during SICD (Avtukh et al. 2023).

Summarizing all of the above, it can be stated that such a phenomenon as SICD is the result of an imbalance in the cellular pathways of glucose catabolism, which can occur when yeast is incubated with glucose in the absence of other vital nutrients.

## Supporting information

However, in 2023, the proteome epistatic database (https://y5k.bio.ed.ac.uk) became available (Messner et al. 2023) which revealed a very interesting

However, in 2023, the proteome epistatic database (https://y5k.bio.ed.ac.uk) became available (Messner et al. 2023) which revealed a very interesting

## References

Avtukh A, Baskunov B, Keshelava V, Valiakhmetov A (2023) Sugar-Induced Cell Death in the Yeast S. cerevisiae Is Accompanied by the Release of Octanoic Acid, Which Does Not Originate from the Fatty Acid Synthesis Type II Mitochondrial System. In: Applied Microbiology, pp 722–734

Bidiuk VA, Alexandrov AI, Valiakhmetov AY (2021) Extracellular pH and high concentration of potassium regulate the primary necrosis in the yeast Saccharomyces cerevisiae. Arch Microbiol 204:35 doi: 10.1007/s00203-021-02708-6

Bonini BM, Van Dijck P, Thevelein JM (2003) Uncoupling of the glucose growth defect and the deregulation of glycolysis in Saccharomyces cerevisiae tps1 mutants expressing trehalose-6-phosphate-insensitive hexokinase from Schizosaccharomyces pombe. Biochimica et Biophysica Acta (BBA) - Bioenergetics 1606:83–93 doi: 10.1016/S0005-2728(03)00086-0

Chen A, Vargas-Smith J, Tapia H, Gibney PA (2022) Characterizing phenotypic diversity of trehalose biosynthesis mutants in multiple wild strains of Saccharomyces cerevisiae. G3 (Bethesda) 12 doi: 10.1093/g3journal/jkac196

D’Autréaux B, Toledano MB (2007) ROS as signaling molecules: mechanisms that generate specificity in ROS homeostasis. Nature Reviews Molecular Cell Biology 8:813–824 doi: 10.1038/nrm2256

de Alteriis E, Cartenì F, Parascandola P, Serpa J, Mazzoleni S (2018) Revisiting the Crabtree/Warburg effect in a dynamic perspective: a fitness advantage against sugar-induced cell death. Cell Cycle 17:688–701 doi: 10.1080/15384101.2018.1442622

De Silva-Udawatta MN, Cannon JF (2001) Roles of trehalose phosphate synthase in yeast glycogen metabolism and sporulation. Molecular Microbiology 40:1345–1356 doi: 10.1046/j.1365-2958.2001.02477.x

François J, Parrou JL (2001) Reserve carbohydrates metabolism in the yeast Saccharomyces cerevisiae. FEMS Microbiol Rev 25:125–145 doi: 10.1111/j.1574-6976.2001.tb00574.x

Gancedo C, Flores C-L (2004) The importance of a functional trehalose biosynthetic pathway for the life of yeasts and fungi. FEMS Yeast Research 4:351–359

Gibney PA et al. (2020) A tps1Δ persister-like state in Saccharomyces cerevisiae is regulated by MKT1. PLoS One 15:e0233779 doi: 10.1371/journal.pone.0233779

Granot D, Dai N (1997) Sugar induced cell death in yeast is dependent on the rate of sugar phosphorylation as determined by Arabidopsis thaliana hexokinase. Cell Death Differ 4:555–559 doi: 10.1038/sj.cdd.4400280

Granot D, Levine A, Dor-Hefetz E (2003) Sugar-induced apoptosis in yeast cells. FEMS Yeast Res 4:7–13

Granot D, Snyder M (1991) Glucose induces cAMP-independent growth-related changes in stationary-phase cells of Saccharomyces cerevisiae. Proc Natl Acad Sci U S A 88:5724–5728

Granot D, Snyder M (1993) Carbon source induces growth of stationary phase yeast cells, independent of carbon source metabolism. Yeast 9:465–479 doi: 10.1002/yea.320090503

Hohmann S, Bell W, NevesA MJ, Valckx D, Thevelein JM (1996) Evidence for trehaloseL6LphosphateLdependent andLindependent mechanisms in the control of sugar influx into yeast glycolysis. Molecular microbiology 20:981–991

Hohmann S, Neves MJ, de Koning W, Alijo R, Ramos J, Thevelein JM (1993) The growth and signaling defects of the ggs1 (fdp1/byp1) deletion mutant on glucose are suppressed by a deletion of the gene encoding hexokinase PII. Current Genetics 23:281–289 doi: 10.1007/BF00310888

Hohmann S, Van Dijck P, Luyten K, Thevelein JM (1994) The byp1-3 allele of the Saccharomyces cerevisiae GGS1/TPS1 gene and its multi-copy suppressor tRNAGLN (CAG): Ggs1/Tps1 protein levels restraining growth on fermentable sugars and trehalose accumulation. Current Genetics 26:295–301 doi: 10.1007/BF00310492

Hottiger T, Schmutz P, Wiemken A (1987) Heat-induced accumulation and futile cycling of trehalose in Saccharomyces cerevisiae. Journal of Bacteriology 169:5518–5522 doi: 10.1128/jb.169.12.5518-5522.1987

Kalyanaraman B et al. (2012) Measuring reactive oxygen and nitrogen species with fluorescent probes: challenges and limitations. Free radical biology & medicine 52:1–6 doi: 10.1016/j.freeradbiomed.2011.09.030

Lee YJ et al. (2011) Phosphate and succinate use different mechanisms to inhibit sugar-induced cell death in yeast: insight into the Crabtree effect. J Biol Chem 286:20267–20274 doi: 10.1074/jbc.M110.209379

Lillie SH, Pringle JR (1980) Reserve carbohydrate metabolism in Saccharomyces cerevisiae: responses to nutrient limitation. Journal of Bacteriology 143:1384–1394 doi: 10.1128/jb.143.3.1384-1394.1980

Liu Y, Wood NE, Marchand AJ, Arguello-Miranda O, Doncic A (2020) Functional interrelationships between carbohydrate and lipid storage, and mitochondrial activity during sporulation in Saccharomyces cerevisiae. Yeast 37:269–279 doi: 10.1002/yea.3460

Lobo Z, Maitra PK (1983) Phosphofructokinase mutants of yeast. Biochemistry and genetics. J Biol Chem 258:1444–1449

Messner CB et al. (2023) The proteomic landscape of genome-wide genetic perturbations. Cell 186:2018-2034.e2021 doi: 10.1016/j.cell.2023.03.026

Murphy Michael P et al. (2011) Unraveling the Biological Roles of Reactive Oxygen Species. Cell Metabolism 13:361–366 doi: 10.1016/j.cmet.2011.03.010

Neves MJ et al. (1995) Control of glucose influx into glycolysis and pleiotropic effects studied in different isogenic sets of Saccharomyces cerevisiae mutants in trehalose biosynthesis. Current Genetics 27:110–122 doi: 10.1007/BF00313424

Parrou JL, Teste M-A, François J (1997) Effects of various types of stress on the metabolism of reserve carbohydrates in Saccharomyces cerevisiae: genetic evidence for a stress-induced recycling of glycogen and trehalose. Microbiology 143:1891–1900 doi: 10.1099/00221287-143-6-1891

Paul MJ, Primavesi LF, Jhurreea D, Zhang Y (2008) Trehalose metabolism and signaling. Annu Rev Plant Biol 59:417–441 doi: 10.1146/annurev.arplant.59.032607.092945

Pereira MD, Eleutherio ECA, Panek AD (2001) Acquisition of tolerance against oxidative damage in Saccharomyces cerevisiae. BMC Microbiology 1:11 doi: 10.1186/1471-2180-1-11

Singer MA, Lindquist S (1998a) Multiple Effects of Trehalose on Protein Folding In Vitro and In Vivo. Molecular Cell 1:639–648 doi: 10.1016/S1097-2765(00)80064-7

Singer MA, Lindquist S (1998b) Thermotolerance in Saccharomyces cerevisiae: the Yin and Yang of trehalose. Trends in Biotechnology 16:460–468 doi: 10.1016/S0167-7799(98)01251-7

Tapia H, Young L, Fox D, Bertozzi CR, Koshland D (2015) Increasing intracellular trehalose is sufficient to confer desiccation tolerance to Saccharomyces cerevisiae. Proceedings of the National Academy of Sciences 112:6122–6127 doi: 10.1073/pnas.1506415112

Thevelein JM, Hohmann S (1995) Trehalose synthase: guard to the gate of glycolysis in yeast? Trends Biochem Sci 20:3–10 doi: 10.1016/s0968-0004(00)88938-0

Valiakhmetov AY, Kuchin AV, Suzina NE, Zvonarev AN, Shepelyakovskaya AO (2019) Glucose causes primary necrosis in exponentially grown yeast Saccharomyces cerevisiae. FEMS Yeast Res 19 doi: 10.1093/femsyr/foz019

van Heerden JH et al. (2014) Lost in transition: start-up of glycolysis yields subpopulations of nongrowing cells. Science 343:1245114 doi: 10.1126/science.1245114

Van Leemputte F, Vanthienen W, Wijnants S, Van Zeebroeck G, Thevelein JM (2020) Aberrant intracellular pH regulation limiting glyceraldehyde-3-phosphate dehydrogenase activity in the glucose-sensitive yeast tps1 Δ mutant. Mbio 11:10.1128/mbio.02199-02120

Walther T et al. (2013) Metabolic phenotypes of Saccharomyces cerevisiae mutants with altered trehalose 6-phosphate dynamics. The Biochemical journal 454:227–237 doi: 10.1042/BJ20130587

Wilson RA, Jenkinson JM, Gibson RP, Littlechild JA, Wang ZY, Talbot NJ (2007) Tps1 regulates the pentose phosphate pathway, nitrogen metabolism, and fungal virulence. The EMBO Journal 26:3673–3685 doi: 10.1038/sj.emboj.7601795

Yoshimoto H et al. (2009) Sugar induces death of the bottom-fermenting yeast Saccharomyces pastorianus. J Biosci Bioeng 108:60–62 doi: 10.1016/j.jbiosc.2008.12.022

