## Supplementary figures and images for "Suppression of glycolysis decreases sugar-induced cell death in Saccharomyces cerevisiae"

### However, in 2023, the proteome epistatic database (https://y5k.bio.ed.ac.uk) became available (Messner et al. 2023) which revealed a very interesting

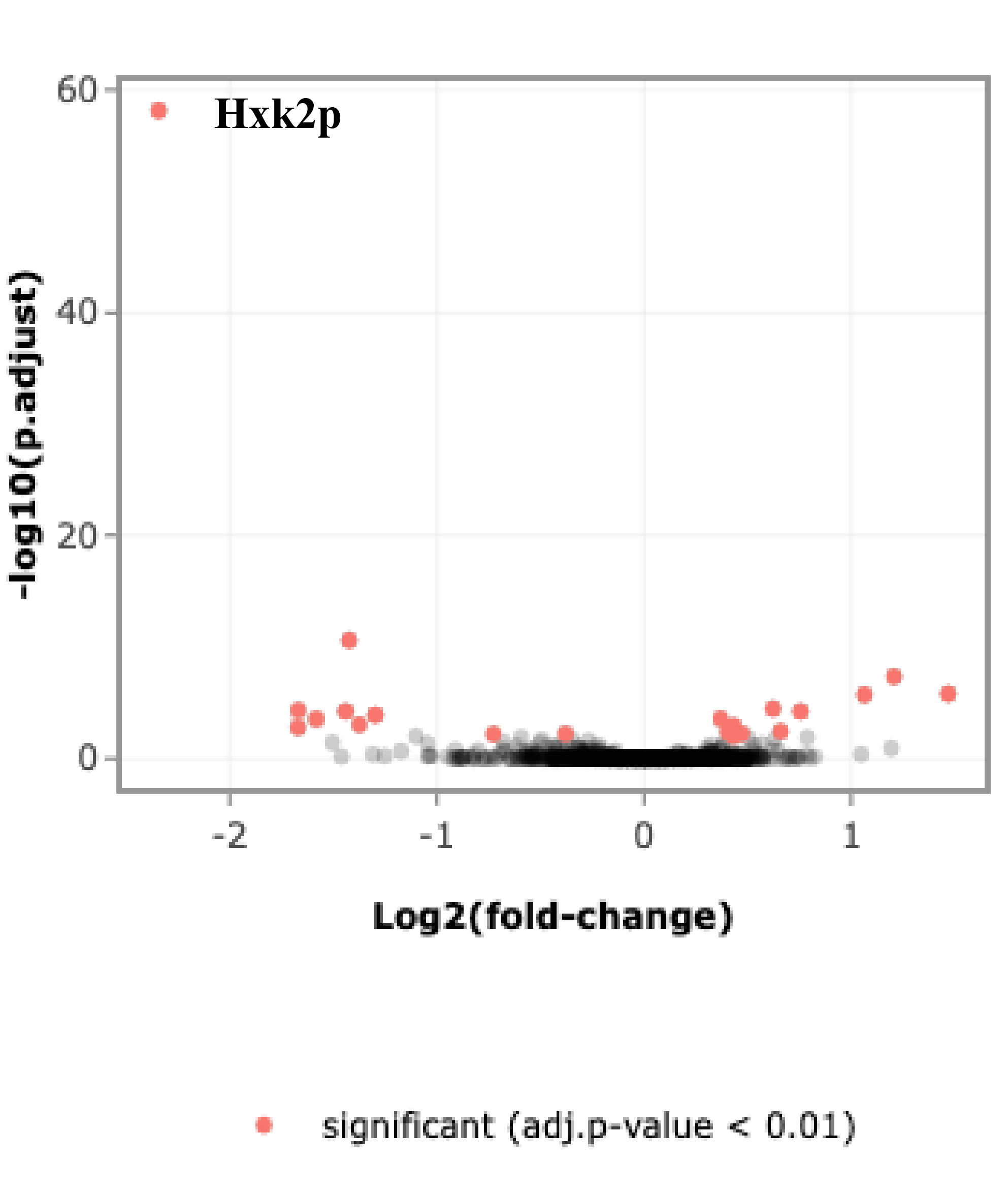
